# Whole-Brain Cell-Cell Interaction Axes Explaining Tissue Vulnerability Across the Neurodegenerative Spectrum

**DOI:** 10.1101/2025.07.21.665217

**Authors:** Veronika Pak, Joon Hwan Hong, Tobias Robert Baumeister, Gleb Bezgin, Corina Nagy, Simon Ducharme, Mahsa Dadar, Yashar Zeighami, Yasser Iturria-Medina

**Author notes:** Correspondence to: Yasser Iturria Medina, PhD, Montreal Neurological Institute, McGill University, Quebec, Canada, H3A 2B4.

## Abstract

Disrupted cell-cell communication represents a fundamental mechanism underlying neurodegeneration, yet how specific intercellular signaling patterns relate to regional brain vulnerability remains poorly understood. Here, we map whole-brain intercellular interaction networks and their spatial correspondence with tissue damage across 13 neurodegenerative conditions, including early– and late-onset Alzheimer’s disease, presenilin-1 mutations, clinical and pathological subtypes of frontotemporal lobar degeneration, Parkinson’s disease, dementia with Lewy bodies, and amyotrophic lateral sclerosis. By integrating multiregional single-nucleus and bulk RNA-seq data with curated cell-cell interaction databases and structural MRI, we reconstruct over 1,000 whole-brain maps of ligand-receptor interactions and quantify their associations with regional atrophy patterns. Multivariate analysis identifies three dominant axes of intercellular communication that explain regional vulnerability to neurodegeneration. Notably, the first axis involves neuron-astrocyte-microglia interactions, explaining atrophy patterns shared by frontotemporal lobar degeneration and Alzheimer’s disease subtypes. Two complementary axes involving neurons, endothelial cells, and astrocytes explain patterns specific to mutations in PS1 and Parkinson’s disease. Importantly, validation in an independent post-mortem cohort (N = 375) confirms that late-onset Alzheimer’s disease-associated cell-cell interactions predict observed frontal cortex atrophy. These results establish a systematic framework linking local intercellular communication networks to spatial patterns of neurodegeneration, revealing both shared and disease-specific molecular pathways that drive regional brain vulnerability and identifying cellular interaction targets for precision therapeutic interventions.

## Introduction

A fundamental question in neuroscience concerns how distinct brain cell populations contribute to neurodegenerative disease pathogenesis (Balusu et al., 2023; Green et al., 2024). While traditionally viewed through a neuron-centric lens (Gorman, 2008), compelling evidence now indicates that non-neuronal cells serve as critical drivers of neurodegeneration (Balusu et al., 2023; Brandebura et al., 2023; Gao et al., 2023; Park et al., 2023; Patani et al., 2023). Advanced multi-omics investigations have reinforced the essential role these cells play in neurodegenerative processes (Li et al., 2023; Martirosyan et al., 2024; Mathys et al., 2019). Cell-cell communication represents a particularly important focus, as its precise regulation underpins the nervous system’s equilibrium and health (Su et al., 2024). For instance, dysfunctions in signalling between neurons and glial cells, including microglia, astrocytes, and oligodendrocytes, are thought to lead to inflammation, impaired glutamate metabolism, demyelination, and consequent neuronal death (Chi et al., 2025; Cihankaya et al., 2022; Saavedra et al., 2022). Characterizing how these cellular interactions modulate vulnerability to diverse neurodegenerative conditions is essential for advancing our mechanistic understanding and for the development of novel therapies.

Significant advances in multi-omics, particularly single-cell and single-nucleus RNA sequencing (snRNA-seq) combined with spatial transcriptomics, have enabled studying cell-cell communication in health and disease (Armingol et al., 2024; Armingol et al., 2020). Despite these advances, detecting global patterns in cell communication across the entire human brain is still challenging, as these methods are often limited to isolated regions of interest. In the context of neurodegeneration, understanding whole-brain patterns is essential since the clinical manifestations of different conditions vary greatly depending on the specific anatomical/functional pathways affected (Pandya & Patani, 2021). Therefore, the limited tissue coverage provided by these methods constrains the ability to conduct systematic analyses across different brain regions and tissue damage patterns specific to different diseases.

In this study, we characterize whole-brain patterns of neurotypical cell-cell interactions spatially aligned with tissue damage observed in 13 neurodegenerative conditions, including early– and late-onset AD (EOAD and LOAD), genetic mutations in presenilin-1 (PS1 or PSEN1), Parkinson’s disease (PD), dementia with Lewy bodies (DLB), amyotrophic lateral sclerosis (ALS), and both clinical and pathological subtypes of frontotemporal lobar degeneration (FTLD) (Dadar et al., 2020; Harper et al., 2017; Metz et al., 2025; Zeighami et al., 2015). First, we generate whole-brain cell-type-specific maps of molecular interactions derived from literature-curated databases, using spatial gene expression and magnetic resonance imaging (MRI) data. Next, using multivariate statistical approaches, we identify principal axes of cell-cell interactions underlying spatial vulnerability patterns across the 13 neurodegenerative conditions. We proceed to conduct gene enrichment analyses on key cell-cell interactions from the prominent axes to uncover underlying signalling pathways predisposing atrophy across the neurodegeneration. Finally, we used an independent LOAD dataset (N = 375) with matched brain RNA and atrophy evaluations to validate the key interactions identified with our *in-silico* approach, confirming the generalizability of our findings. This study represents an advancement in our understanding of neurodegenerative diseases, with potential implications for the development of targeted therapeutics that modulate specific cell-cell interactions. Notably, by identifying the cellular communication networks that underlie regional vulnerability, we provide a novel framework for understanding why certain brain regions are primarily targeted by specific neurodegenerative conditions.

## Materials and methods

### Allen Human Brain Atlas

Bulk microarray messenger RNA (mRNA) expression data from the Allen Human Brain Atlas (AHBA; https://human.brain-map.org/) was used to construct interaction maps (Hawrylycz et al., 2012). This included anatomical and histological data collected post-mortem from six healthy individuals with no history of neurological disease (one female; age range 24-57 years; mean age 42.5 ± 13.38 years) (Hawrylycz et al., 2012). Data from two donors covered the entire brain, while the remaining four included the left hemisphere only. A total of 3,702 spatially distinct samples were distributed across cortical, subcortical, brainstem, and cerebellar regions, with expression levels measured for over 20,000 genes (Hawrylycz et al., 2012). Each brain sample is labeled with native MRI coordinates (Talairach & Szikla, 1980) and Montreal Neurological Institute (MNI) coordinates (Evans et al., 1994), based on structural MRI collected from the same participants, allowing precise alignment of samples with imaging data. Data were initially preprocessed by the Allen Institute to reduce the effects of bias due to batch effects (Hawrylycz et al., 2012).

To estimate gene expression levels at unsampled brain locations, we applied Gaussian process regression, a previously validated method to interpolate AHBA data across the whole brain (Gryglewski et al., 2018). This approach allows the prediction of mRNA expression levels for all gray matter (GM) voxels in the standardized MNI brain space (Evans et al., 1994). Gaussian process regression predicts gene expression in unobserved regions using values from neighboring sampled regions, with weights decreasing as distance increases. For genes represented by multiple microarray probes, this framework was additionally used to evaluate probe-specific spatial predictability via leave-one-out cross-validation, and the probe with the highest prediction accuracy was selected as the representative probe. Interpolation was performed on pooled multi-donor samples.

### Harvard Brain Tissue Resource Center

Adjusted gene expression data from autopsied dorsolateral prefrontal cortex (DLPFC) brain tissue of LOAD patients were obtained from the Gene Expression Omnibus (GEO) database under accession number GSE44770. This dataset included detailed demographic information, disease states, and measurements of brain atrophy (Zhang et al., 2013). Brain tissue samples were originally collected through the Harvard Brain Tissue Resource Center (HBTRC), where all subjects underwent a comprehensive diagnostic assessment upon intake, and each brain was rigorously examined for LOAD-related neuropathology (Zhang et al., 2013). Gene expression levels had been previously adjusted to account for covariates, including age, sex, postmortem interval (PMI) in hours, sample pH, and RNA integrity number (RIN) (Zhang et al., 2013). Of the initial 379 participants with AD diagnosis, 375 were included in our analysis (mean age 80.81 ± 9.05 years; 156 males, 219 females), as complete data for both gene expression and corresponding regional atrophy in the prefrontal cortex were available for these individuals.

### Allen Brain Cell Atlas

The Allen Brain Cell (ABC) Atlas is a platform that explores single-cell data across the mammalian brain. In this study, we used an ABC dataset profiling over 3 million cells from the entire adult human brain, generated using snRNA-seq (10x Genomics) (Siletti et al., 2023). Post-mortem brain tissue was collected from three neurotypical male donors (ages 29-60; mean age 40.33 ± 10.60 years) with no known history of neuropsychiatric or neurological conditions (Siletti et al., 2023). One additional motor cortex dissection was included from a 60-year-old female donor (Siletti et al., 2023). The dataset includes 109 standardized dissections spanning the cerebral cortex and subcortical regions (Siletti et al., 2023). Cells were hierarchically clustered into 31 superclusters, 461 clusters, and 3,313 subclusters (Siletti et al., 2023). Here, we leveraged the ABC atlas to independently validate region-specific cell-cell interactions identified using bulk transcriptomic data from AHBA (Siletti et al., 2023).

### Ligand-receptor interaction databases

Databases of interaction pairs were obtained from the “Compendium of Available Lists of Ligand-Receptor Pairs in Literature” curated by Armingol et al. in their systematic review of cell-cell communication inference methods (Armingol et al., 2021). Ten databases of LR pairs specific to human tissues were selected (Browaeys et al., 2020; Cabello-Aguilar et al., 2020; Dimitrov et al., 2022; Hou et al., 2020; Jin et al., 2021; Noel et al., 2021; Ramilowski et al., 2015; Shao et al., 2021; Wang et al., 2019), including NeuronChatDB, which focuses on neuronal interactions (Zhao et al., 2023). Ligand-receptor (LR) pairs involving composite multi-gene complexes or cofactors were mostly excluded to maintain consistent one-to-one gene pair formatting. After concatenating all pairs across databases, we compiled a unique list of genes corresponding to ligands and receptors. This list was cross-referenced with the AHBA database of approximately 20,000 genes, resulting in 5,511 LR pairs available.

### Cell type annotation

To capture interactions only relevant to the human brain, we limited our analysis to genes corresponding to ligands and receptors frequently expressed in six major brain cell types: neurons, astrocytes, microglia, endothelial cells, oligodendrocytes, and oligodendrocyte precursor cells. We developed an R package BrainCellAnn, which integrates comprehensive reference databases of gene markers specific to cell types in the human brain, achieving consensus across numerous multi-omics datasets.

The package estimates consensus cell types through comparing outputs from different databases, ensuring specificity and reliability. The first database used was the R package BRETIGEA, which includes brain gene markers from five RNA-seq studies, validated by ATAC-seq and immunohistochemistry analyses on the same brain samples (McKenzie et al., 2018). The second resource was the PanglaoDB database of curated gene signatures, integrated into the OmniPath (Türei et al., 2021) and decoupleR (Badia-i-Mompel et al., 2022) R packages. PanglaoDB contains pre-processed meta-analyses from over 1,054 scRNA-seq experiments, covering a wide range of tissues and organs (Franzén et al., 2019). Thirdly, we used the CellMarker 2.0 database (Hu et al., 2023), which provides a manually curated collection of experimentally supported markers for various cell types in human and mouse tissues, including markers from approximately 102,000 studies in PubMed and cells derived from 48 snRNA-seq studies. The final resources were the immunohistochemistry and snRNA-seq from the Human Protein Atlas (HPA), the largest and most comprehensive web database providing information on protein expression and localization across human tissues and cells (Thul & Lindskog, 2018).

The algorithm of the package takes a list of gene markers as input, then each database returns one or two most likely candidate cell types. For a cell type to reach consensus, it should be labelled in agreement with at least two databases. The consensus score is calculated based on number of appearances of a cell type per database and database-specific weights. In cases of consensus conflict, database-specific weights are ranked using Random Forest feature importance to match cell type annotations from the Allen Institute’s ABC Whole-Brain Human Atlas (Siletti et al., 2023), prioritizing certain databases for specific cell types over others.

Out of 5,511 interaction pairs, only 1,037 had both the ligand and receptor consistently annotated by at least two databases. Considerations for immune cells were made, so genes known to be highly expressed in T cells, B cells, and Natural Killer cells were excluded.

### Reconstruction of interaction maps

Methods for inferring cell-cell communication for snRNA-seq data such as CellPhoneDB (Efremova et al., 2020), Connectome (Raredon et al., 2022), NATMI (Hou et al., 2020), and ICELLNET (Noel et al., 2021) use the product of ligand and receptor expression levels as a measure of communication. Similarly, in the AHBA bulk data we use here, cell-cell communication was calculated by multiplying the expression level of a ligand in an individual voxel (1.5 x 1.5 x 1.5 mm^3^ size) by the expression level of the corresponding receptor in the same voxel:

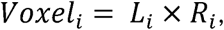

where *i* represents an individual voxel (i = 1, 2, …, n), *L* is the expression level of a ligand in the voxel, and *R* is the expression level of a receptor gene in the voxel. The communication strength reflects co-expression of ligand and receptor genes within mixed cell populations.

Interactions between individual LR pairs were estimated across all gray matter voxels. For each map, interaction measures were registered into MNI and volumetric space using the ICBM152 template (Evans et al., 1994).

For the ABC dataset, gene expression was measured specifically from the cell type corresponding to each annotated by the package ligand or receptor in the 1,037 interaction pairs. For each gene, expression from the matched cell type was first averaged across regions, and then, for each interacting pair, the ligand and receptor regional expression values were multiplied.

### Disorder-specific atrophy maps

We obtained whole-brain voxel-wise atrophy maps for thirteen neurodegenerative conditions, including early– and late-onset Alzheimer’s disease (EOAD and LOAD, respectively), Parkinson’s disease (PD), amyotrophic lateral sclerosis (ALS), dementia with Lewy bodies (DLB), mutations in presenilin-1 (PS-1), clinical variants of frontotemporal dementia (FTD; the behavioural variant bvFTD and the non-fluent and semantic variants of primary progressive aphasia nfvPPA and svPPA), and FTLD-related pathologies such as FLTD-TDP (TAR DNA-binding protein) types A and C, 3-repeat tauopathy, and 4-repeat tauopathy (3RTau and 4RTau) (Dadar et al., 2020; Harper et al., 2017; Zeighami et al., 2015). The term FTD was used when addressing the clinical syndromes, and the term FTLD when referencing the histologically confirmed neurodegenerative pathologies (Boeve et al., 2022). Changes in tissue density in the atrophy maps were measured by voxel– and deformation-based morphometry (VBM and DBM) applied to structural T1-weighted MR images, and expressed as a t-score per voxel (relatively low negative values indicate greater GM tissue loss/atrophy) (Ashburner & Friston, 2000; Aubert-Broche et al., 2013). VBM is a data-driven technique to identify regional differences in tissue concentration throughout the whole brain, without requiring predefined regions of interest (Ashburner & Friston, 2000). DBM is related method for detecting structural brain changes across participants, which in addition considers anatomical differences such as shape and size of brain structures (Aubert-Broche et al., 2013). **Supplementary Table 1** summarizes origin of each disorder-related atrophy map compiled from different studies. Initially, all atrophy maps except for PD had negative values indicating atrophy and positive values indicating tissue enlargement compared to controls; however, these values were reversed for an easier and more intuitive interpretation of their relationships with LR interactions in the Results section.

MRI data for neuropathological dementias were obtained from 186 individuals with clinically diagnosed dementia, confirmed by histopathological examination (either postmortem or biopsy), along with 73 cognitively healthy controls (Harper et al., 2017). Data were averaged by condition: 107 participants had a primary diagnosis of Alzheimer’s disease (AD) (including 68 with early-onset [<65 years at onset], 29 with late-onset [≥65 years at onset], and 10 carriers of the PS1 mutation), 25 had dementia with Lewy bodies (DLB), 11 had three-repeat tauopathy, 17 had four-repeat tauopathy, 12 had frontotemporal lobar degeneration with TDP-43 type A (FTLD-TDP type A), and 14 had FTLD-TDP type C (Harper et al., 2017). T1-weighted volumetric MRI data were collected from participants during life across multiple centers, using scanners from three manufacturers (Philips, GE, and Siemens) with different imaging protocols (Harper et al., 2017). Magnetic field strengths included 1.0 T (15 scans), 1.5 T (201 scans), and 3 T (43 scans) (Harper et al., 2017). Pathological examination of brain tissue took place between 1997 and 2015, following standard histopathological procedures and criteria at one of four centers: the Queen Square Brain Bank in London, Kings College Hospital in London, VU Medical Centre in Amsterdam, and the Institute for Ageing and Health in Newcastle (Harper et al., 2017). Atrophy maps were statistically adjusted for age, sex, total intracranial volume, and MRI field strength and site (Harper et al., 2017). Ethical approval for the corresponding study was provided by the National Research Ethics Service Committee London-Southeast (Harper et al., 2017).

For PD, MRI data were obtained from the Parkinson’s Progression Markers Initiative (PPMI) database (Parkinson Progression Marker, 2011; Zeighami et al., 2015). The PPMI is a multicenter international study with acquisition protocols approved by the local institutional review boards at all 24 sites across the United States, Europe, and Australia (Parkinson Progression Marker, 2011). Imaging data were collected at 16 centers participating in the PPMI project, using scanners from three different manufacturers (GE Medical Systems, Siemens, and Philips Medical Systems) (Parkinson Progression Marker, 2011). High-resolution 3T T1-weighted MRI scans from the initial visit, along with clinical data used to construct atrophy maps, were acquired from 232 participants with PD and 118 age-matched controls (Zeighami et al., 2015). PD subjects (77 females; age 61.2 ± 9.1) were required to be at least 30 years old, untreated with PD medications, diagnosed within the last two years, and exhibit at least two PD-related motor symptoms, such as asymmetrical resting tremor, uneven bradykinesia, or a combination of bradykinesia, resting tremor, and rigidity (Parkinson Progression Marker, 2011). All participants underwent dopamine transporter (DAT) imaging to confirm a DAT deficit as a prerequisite for eligibility (Parkinson Progression Marker, 2011). Written informed consent was collected from all study participants (Parkinson Progression Marker, 2011).

MRI data from ALS patients were obtained from 66 individuals (24 females; age 57.98 ± 10.84), as well as 43 matched control individuals, who participated in the longitudinal study through the Canadian ALS Neuroimaging Consortium (ClinicalTrials.gov NCT02405182) (Dadar et al., 2020; Kalra et al., 2020). This consortium includes 3T MRI sites at the University of Alberta, University of Calgary, University of Toronto, and McGill University (Kalra et al., 2020). Patients were eligible if they were diagnosed with sporadic or familial ALS and met the revised research criteria from El Escorial (Brooks et al., 2000) for possible, laboratory-supported, or definite ALS (Kalra et al., 2020). Neurological examination was conducted by a trained neurologist at each participating site (Kalra et al., 2020). Clinical evaluations included global measures of disease status, upper motor neuron dysfunction and cognitive batteries (Kalra et al., 2020). Written informed consent was obtained from all participants, and the study received ethical approval from the health research ethics boards at each center (Kalra et al., 2020). Exclusion criteria included a history of other neurological or psychiatric disorders, prior brain injury, or respiratory impairment that prevented tolerance of the MRI protocol (Kalra et al., 2020). Patients with primary lateral sclerosis, progressive muscular atrophy, or FTD were also excluded (Kalra et al., 2020). Before constructing the maps, normative aging and sex differences were regressed out of the data (Dadar et al., 2020).

For clinical subtypes of FTD, data were examined from the Frontotemporal Lobar Degeneration Neuroimaging Initiative (FTLDNI AG032306), part of the ALLFTD initiative. Atrophy maps were constructed using data from 136 FTD patients and 133 age-matched control participants (Metz et al., 2025). Participants were grouped according to their FTD clinical variant: 70 were diagnosed with the behavioral variant, 36 with semantic primary progressive aphasia, and 30 with non-fluent primary progressive aphasia (Metz et al., 2025; Staffaroni et al., 2019). Structural 3T MRI scans were acquired at three locations: University of California San Francisco, Mayo Clinic, and Massachusetts General Hospital (Staffaroni et al., 2019). Patients were either referred by physicians or self-referred and underwent comprehensive neurological, neuropsychological, and functional assessments, including informant interviews (Staffaroni et al., 2019). Diagnoses were established during multidisciplinary consensus conferences based on either the Neary criteria (Neary et al., 1998) or, depending on enrollment year, the more recent consensus criteria for bvFTD (Rascovsky et al., 2011) and PPA (Gorno-Tempini et al., 2011). Histological analysis was also conducted to assess potential AD pathology, due to overlapping clinical symptoms between the conditions (Staffaroni et al., 2019). All participants provided informed consent, and the study protocol received institutional review board approval at each site (Staffaroni et al., 2019).

### Anatomical parcellations

Two anatomical parcellations were used to define regions of interest in the validation analyses. The Desikan-Killiany-Tourville (DKT) parcellation included 62 distinct cortical regions, which were averaged across both left and right hemispheres (Desikan et al., 2006; Klein & Tourville, 2012). We then manually added several subcortical structures from FreeSurfer segmentation, including the subthalamic nucleus, red nucleus, dentate nucleus, and substantia nigra, resulting in 104 (43 when averaged across hemispheres and cerebellar structures) unique regions used for validating dopamine-related interaction maps.

The Allen Human Reference Atlas included 141 brain regions from both hemispheres of the adult human brain (Ding et al., 2016). Many of these regions corresponded to modified Brodmann areas and were designed to reflect cellular resolution in histological images, helping to link microscopic features with the macroscopic scales common in neuroimaging studies (Ding et al., 2016). This atlas was also used to define many of the regions included in the ABC atlas (Siletti et al., 2023), which we used to validate our bulk-derived cell-cell interactions. In total, we used 76 brain regions that overlapped between the ABC atlas and the Allen Human Reference Atlas and were available in our cell-cell interaction maps.

### Spatial correlation with PET-measured dopamine

To examine spatial correlation between molecular interaction maps and dopamine receptor/transporter availability, we used the JuSpace toolbox (Dukart et al., 2021) to correlate regional interaction values with positron emission tomography (PET) and single-photon emission computed tomography (SPECT) imaging data. D1 receptor-selective [^11^C]SCH23390 maps were measured by Kaller et al. (2017), [^18^F]Fallypride D2/3 receptor binding maps were obtained from Jaworska et al. (2020), and dopamine transporter (DAT) availability maps were obtained from Dukart et al. (2018).

Prior to correlation, both molecular interaction and imaging data were rank-transformed, and spatial associations were quantified using Spearman’s rank correlation coefficient (ρ). Correlation coefficients were Fisher z-transformed to normalize their distribution. To account for spatial autocorrelation, we computed partial correlations adjusting for underlying gray matter probability using the tissue probability map (TPM.nii) from SPM12. Statistical significance was assessed using permutation testing with 1,000 iterations.

### Partial least squares analyses

Multidimensional associations between atrophy patterns specific to 13 neurodegenerative conditions and 1,037 LR interaction scores were uncovered using partial least squares (PLS) cross-correlation analysis (Geladi & Kowalski, 1986; Lin et al., 2022; McIntosh & Lobaugh, 2004). PLS correlation, a data reduction technique related to principal component analysis (PCA), uses singular value decomposition (SVD) on the covariance matrix of two datasets (X and Y) to identify linear combinations (latent variables) of the original variables that optimally covary (Geladi & Kowalski, 1986; McIntosh & Lobaugh, 2004).

Mathematically, the SVD of the covariance matrix is expressed as:

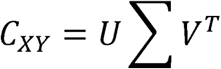

where U and V represent orthogonal matrices containing the singular vectors, and ∑ is a diagonal matrix containing the singular values indicating the strength of covariation.

We applied PLS correlation to rank interactions based on their multivariate spatial alignment with atrophy patterns across 13 neurodegenerative conditions. Prior to PLS analysis, all values were z-score normalized. Unlike PCA, which maximizes variance within a single dataset (X), PLS maximizes the covariance between two datasets (X and Y), where X represents interactions, and Y represents atrophy patterns.

Specifically, PLS seeks weight vectors w_X_ and w_Y_ to maximize:

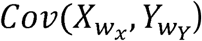

The resulting weights and factor scores ensure maximum covariance, facilitating the identification of molecular interactions most closely aligned with disease-specific atrophy patterns. To assess the statistical significance of each PLS-derived latent variable or axis, random permutation testing and bootstrapping were performed with 1,000 iterations each. Permutation testing provided family-wise error (FWE) corrected p-values to generate a null distribution of singular values for each axis, while bootstrapping was used to estimate the robustness of feature contribution of each original variable, where 95% confidence intervals (CI) were calculated for each LR interaction/condition feature. The top 10% (104 of the 1,037 interactions) with the largest absolute weights were selected for downstream analysis and interpretation.

### Statistical analyses

We assessed the correspondence between bulk– and single-nucleus-derived LR interactions using partial Spearman correlations, controlling for regional differences. Statistical significance was determined using 500 permutations and resulting p-values were adjusted for multiple comparisons with the false discovery rate (FDR) correction.

To investigate disorder-related commonalities, we applied the K-Means clustering algorithm to the normalized PLS loadings of the 13 neurodegenerative conditions. The Silhouette criterion was used to identify the optimal number of clusters.

### Enrichment analysis with PANTHER

The PANTHER database was used to identify enriched pathways specific to cell signalling that associate with the genes corresponding to top 10% ligands and receptors (Mi & Thomas, 2009). Gene-set enrichment analysis was conducted using the ShinyGO web tool (Ge et al., 2020). The enrichment results were reported as −10log(p-value), with an FDR cutoff of 0.01 applied to control for multiple testing. This cutoff ensured that only highly significant pathways were reported in the results.

## Results

### Reconstructing brain maps of cell-cell interactions

In a first of its kind analysis, we reconstructed whole-brain maps of molecular interactions across six major cell types of the neurotypical human brain (see Fig. 1), including neurons, astrocytes, microglia, endothelial cells, oligodendrocytes, and oligodendrocyte precursor cells (OPCs). First, we utilized bulk microarray gene expression data from six human donors from the AHBA (Gryglewski et al., 2018; Hawrylycz et al., 2012). Next, we integrated 10 curated databases that included literature-supported LR or molecular interaction pairs specific to humans (Armingol et al., 2021; Ramilowski et al., 2015) (see *Methods*, *Ligand-receptor interaction databases*). From the overlap of these 10 databases with the genes available in the AHBA, we identified 1,037 pairs whose ligands and receptors were commonly found to be expressed in the six studied cell types. For a pair to be selected, both the ligand and receptor (or molecule) had to be annotated as being expressed in one of the targeted cell types, with agreement from at least two out of five public gene marker databases (Fig. 1b; see *Methods, Cell type annotation*). The databases included CellMarker 2.0 (Hu et al., 2023), PanglaoDB (Franzén et al., 2019), BRETIGEA (McKenzie et al., 2018), and the immunohistochemistry and single cell databases from the Human Protein Atlas (Thul & Lindskog, 2018).

**Figure 1.**
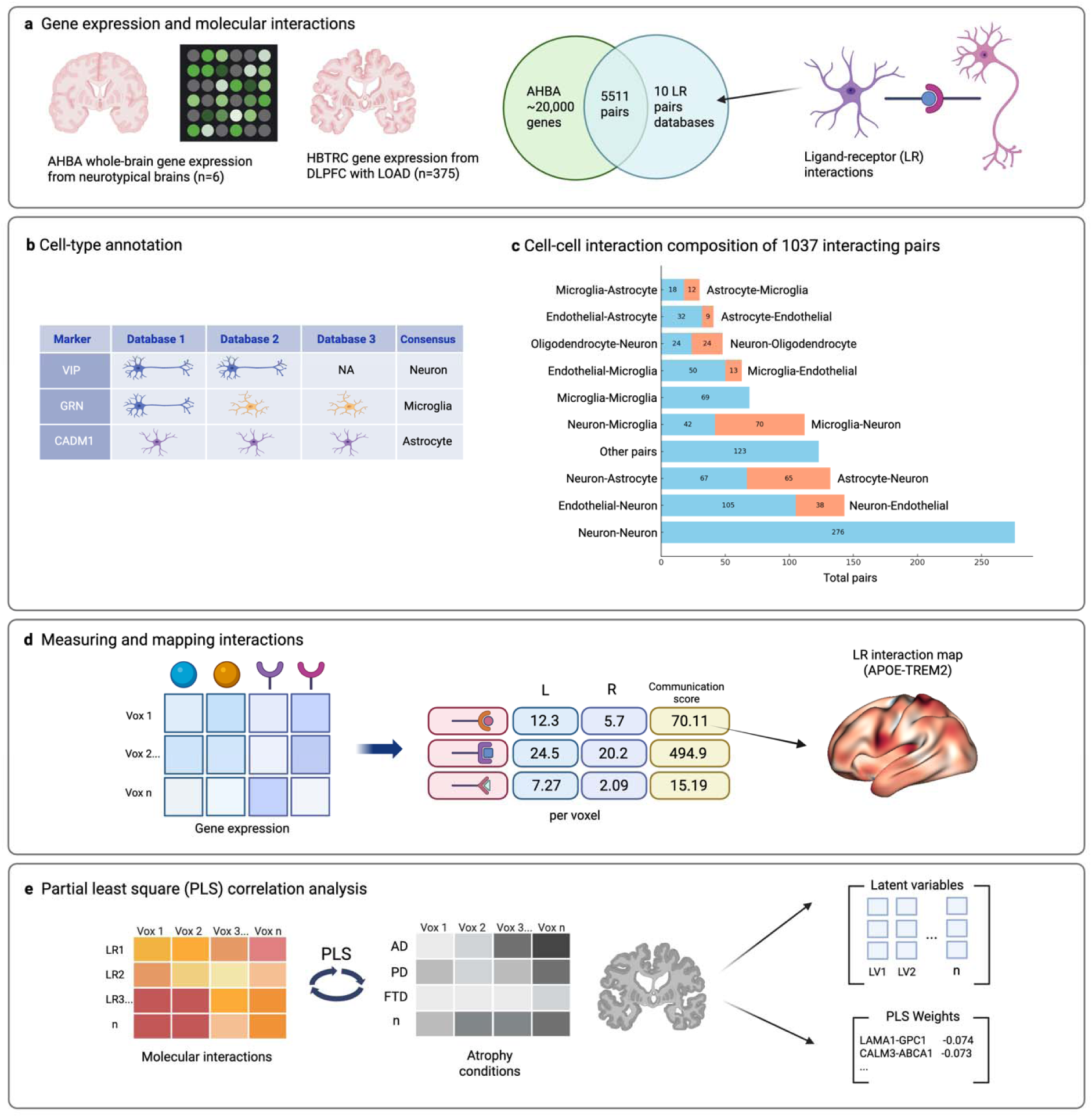
|The process of generating molecular interaction maps and analyzing their relationship with atrophy in neurodegeneration. (**a**) Interpolated whole-brain RNA expression from healthy individuals (Allen Human Brain Atlas; AHBA) and dorsolateral prefrontal cortex (DLPFC) expression from LOAD patients (Harvard Brain Tissue Resource Centre; HBTRC) were analyzed. Interaction pairs from 10 curated human-specific databases (blue circle) were cross-referenced with ∼20,000 genes from the AHBA (green circle). (**b**) The R package BrainCellAnn was used to annotate marker genes associated with interacting pairs. Only pairs whose ligand and receptor genes were annotated as a cell type of interest by at least two marker databases were included in the analysis, resulting in 1,037 pairs. (**c**) Breakdown of the annotated cell-cell interaction ratios, split by direction of interaction types. (**d**) The product of ligand and receptor (or molecule 1 and molecule 2) expression levels within each MNI template voxel (1.5 x 1.5 x 1.5 mm^3^ size) was calculated to generate a whole-brain interaction map for each LR pair. (**e**) PLS correlation was performed at the voxel level, with the 1,037 interaction maps as predictors and atrophy maps for 13 neurodegenerative conditions as outcomes.

The distribution of the different types of annotated cell-cell communication is visualized in Figure 1c. The majority of interactions involved neuron-neuron and neuron-endothelial/endothelial-neuron communication, covering 276 and 143 interacting pairs, respectively. The full list of 1,037 interacting pairs and their cell type annotation is listed in the **Supplementary Table 2**. To quantify brain-wide cell-cell communication for each of the pairs within each voxel of the brain defined by the MNI-152 space (Evans et al., 1994), we calculated the product of ligand and receptor expression levels within each voxel (Fig.1d; see *Methods*, *Reconstruction of interaction maps*). Although most interactions represent direct intercellular LR communication, some non-canonical pairs were also included in the LR databases: receptor complexes, synaptic vesicle proteins, and intracellular signaling components.

Next, to uncover spatial associations between the interaction maps and the atrophy patterns observed in 13 neurodegenerative conditions, we performed partial least squares (PLS) analysis with cross-correlation, using voxels as observations (Fig. 1e), and interactions and disorders as predictor and response variables, respectively. Atrophy maps for the following neurodegenerative conditions were included: early– and late-onset Alzheimer’s disease (EOAD and LOAD, respectively), Parkinson’s disease (PD), amyotrophic lateral sclerosis (ALS), dementia with Lewy bodies (DLB), mutations in presenilin-1 (PS1), clinical variants of FTD (the behavioral variant [bvFTD] and the non-fluent and semantic variants of primary progressive aphasia [nfvPPA and svPPA]), and FTLD-related pathologies such as FLTD-TDP (TAR DNA-binding protein) types A and C, three-repeat tauopathy and four-repeat tauopathy (3RTau and 4RTau)(Dadar et al., 2020; Harper et al., 2017; Metz et al., 2025; Zeighami et al., 2015). The latent variables (hereby axes) identified by PLS, along with their contributing weights, were subsequently analyzed in downstream analyses. In sum, this approach identified the specific molecular interactions and neurodegenerative conditions that contributed most to the covariance between cell-cell communication and tissue atrophy, reflecting potential vulnerabilities.

### Obtained maps of dopamine signalling interactions reflect major dopaminergic pathways

To validate our approach for reconstructing cell-cell interaction maps, we proceeded to test whether the inferred dopamine-related interactions were particularly enriched in key regions of major dopaminergic pathways (e.g., substantia nigra and nucleus accumbens). We selected eight maps of known dopamine interactions that were available in our list, including dopamine receptor families (DRD 1-5) interacting with their corresponding guanine (G) nucleotide-binding proteins and interactions involving the solute carrier family 18-member 2 (*SLC18A2*). SLC18A2 is essential for enabling the release of neurotransmitters, such as dopamine, norepinephrine, serotonin, and histamine, via synaptic vesicles into the synaptic cleft (Surratt et al., 1993). We extracted regional values from the maps defined by the extended DKT atlas (Desikan et al., 2006; Klein & Tourville, 2012), with manually added subcortical regions. **Figure 2a** depicts a heatmap of the regional communication values, which were scaled relative to each other for each interaction.

**Figure 2.**
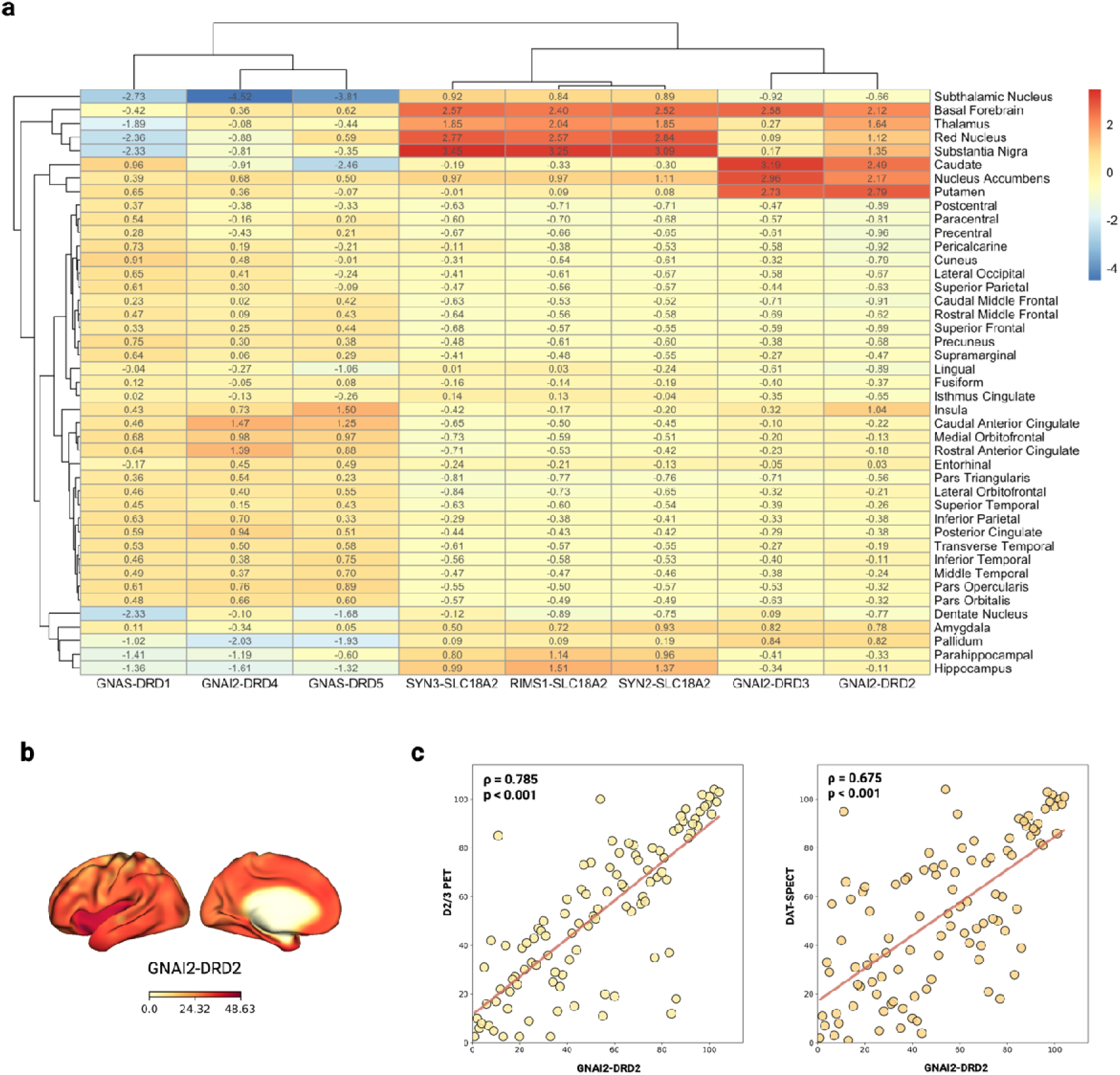
| Regional communication values for dopamine-related interactions. Regions were defined by the extended DKT atlas, with additional subcortical areas included. For each interaction, regional values were normalized, with higher positive scores (in red) indicating stronger communication relative to other regions, and lower negative scores (in blue) representing weaker communication. Regions involved in dopaminergic pathways such as the basal forebrain grouped together and showed higher communication scores for GNAI2-DRD2, GNAI2-DRD3, and SLC18A2 interactions.

We verified (Fig. 2a) that subcortical regions associated with dopamine pathways were enriched with interactions involving GNAI2-DRD3, GNAI2-DRD2, and SLC18A2, compared to other brain regions. Specifically, SLC18A2-related interactions were particularly enriched in the basal forebrain, thalamus, red nucleus, and substantia nigra—regions that play a key role in dopaminergic, cholinergic, and glutamatergic neurotransmission, supporting motor and cognitive functions. Furthermore, interactions involving DRD2 and DRD3 receptors were prominent in striatum regions, including the putamen, caudate, and nucleus accumbens. All three regions receive dopaminergic projections from the ventral tegmental area and the substantia nigra. GNAI2-DRD2 interactions were particularly enriched across all regions heavily involved in the dopamine pathways. In contrast, interactions involving DRD1, DRD4, and DRD5 receptors were not enriched in subcortical regions associated with dopamine pathways.

Next, we tested whether these molecular interactions follow the distribution of D1 (Kaller et al., 2017), D2/3 receptors (Jaworska et al., 2020), and presynaptic dopamine transporters (DAT) (Dukart et al., 2018) measured by PET/SPECT imaging. Using the JuSpace toolbox (Dukart et al., 2021), we computed partial Spearman correlations with spatial autocorrelation correction and 1,000 permutations (**Supplementary Table 3)**. GNAI2-DRD2 and GNAI2-DRD3 interactions showed robust correlations with all dopamine maps (p < 0.001). Specifically, GNAI2-DRD2 (Figs 2b & c) exhibited the strongest associations with D2/3 receptors (Mean Fisher’s z = 1.05, p < 0.001) and DAT (Mean Fisher’s z = 0.81, p < 0.001). In contrast, GNAS-DRD1/DRD5 correlated strongly with D1/D2/D3 PET maps (p < 0.005) but not DAT, consistent with the postsynaptic localization of Gs-coupled D1/D5 receptors. No significant associations were observed for GNAI2-DRD4 interactions. Presynaptic vesicular interactions involving SLC18A2 selectively aligned with DAT maps (p < 0.001) but not with dopamine receptor maps, consistent with the localization of vesicular monoamine transporter 2 and DAT at presynaptic dopamine terminals.

### First axis of LR interactions predominantly captures tissue damage in FTLD and AD

We applied multivariate PLS correlation and permutation tests to explore the spatial covariation between 1,037 interaction pairs and atrophy patterns across 13 neurodegenerative conditions. A thousand permutation and bootstrap iterations were performed to estimate statistical significance and reliability (see *Methods*, *Partial least square analysis*). All 13 latent variables (LVs) or axes of covariation were significant (all p < 0.05, corrected via randomized permutations), possibly due to a large number of observations (∼300,000 voxels). Results for the first three axes are summarized in the **Supplementary Table 4**.

The first axis (latent variable 1) explained 84.27% of the covariance between interactions and atrophy patterns. Figure 3a presents the top 5% of LR pairs with the biggest contributions measured by PLS weights in explaining atrophy. Among these, the COL1A1-CD36 interaction and other CD36-associated pairs in the top 10 contributed most to explaining atrophy patterns (Fig. 3a). CD36 is a membrane glycoprotein that binds Aβ as a scavenger receptor and has been linked to neurodegeneration through Aβ clearance, oxidative stress, neurovascular dysfunction, and chronic inflammation in AD (Dobri et al., 2021; Feng et al., 2024; Park et al., 2011).

**Figure 3.**
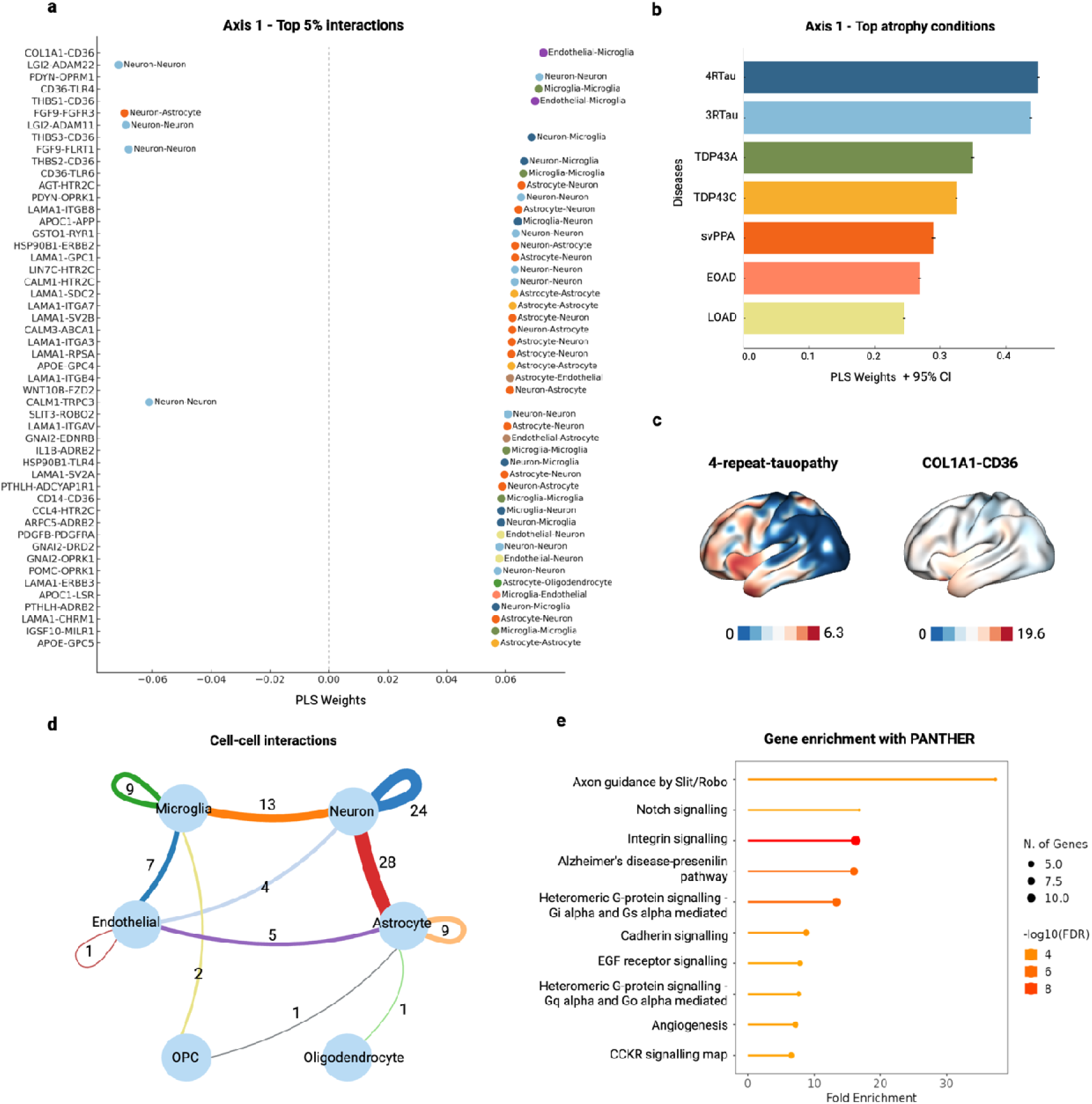
| Top interactions and neurodegenerative conditions contributing to the first axis. (**a**) The top 5% of pairs with the largest contributions to spatial covariance with atrophy are colored by annotated interaction type. The interaction COL1A1-CD36 showed the highest contribution (0.073). PLS weights reflect relative contributions to the axes and are not interpretable as effect sizes or percentages. (**b**) Top neurodegenerative conditions contributing to the first axis, with FTLD-related conditions (4RTau, 3RTau, TDP43A & C, svPPA) making the largest contributions (error bars defined by 95% CI). (**c**) Surface plots (lateral views of the left hemisphere) displaying the atrophy map for 4RTau and the LR interaction map for COL1A1-CD36. Color bars indicate the strength of cell-cell communication and tissue loss, with negative values indicating tissue enlargement relative to healthy controls. (**d**) Network visualization of cell-cell interactions counts from the top 10% annotated pairs, highlighting strong bi-directional signaling between neurons and astrocytes, neurons between each other, and neurons and microglia. Numbers on edges indicate counts of unique interactions. Cell types for interaction types were annotated using five gene marker databases. (**e**) Gene enrichment analysis using the PANTHER database, indicated significant enrichment of pathways related to Slit/Robo-mediated axon guidance, Notch signalling, integrin signalling, and Alzheimer’s disease presenilin (p < 0.001 FDR corrected).

The top 10% (104 interactions) from the first axis included other prominent to neurodegeneration interactions: toll-like receptors (TLR4, TLR6); 12 thrombospondin (THBS) pairs; FGF9-FGFR3, where FGFR3 mediates Aβ-induced tau uptake and neuron-microglia communication; APOE-GPC4, involving the major AD genetic risk factor APOE and astrocyte-secreted GPC4 that drives tau hyperphosphorylation; APOC1-APP, where APOC1 enhances APP cleavage and Aβ production; HSP90B1-ERBB2, with heat shock protein HSP90B1 linked to astrogliosis and tau-driven neurotoxicity; and interactions involving HTR2C (serotonin receptor) and LAMA1 (basal lamina component) (Loch et al., 2023; Saroja et al., 2022; Thümmler et al., 2023; Wei et al., 2024; Zhou et al., 2014).

Figure 3b shows bar graphs representing the atrophy conditions ranked by their contributions to the first axis, with five FTLD-related conditions showing the largest weights, followed by EOAD and LOAD. This suggests that the identified cell-cell interactions mostly explain the atrophy patterns common to the FTLD conditions. Spatial atrophy associated with 4-repeat tauopathy and 3-repeat tauopathy demonstrated the highest contributions, followed by TDP-43 proteinopathy types A and C, and svPPA.

When interpreting these results, a positive association, where both condition and interaction weights are positive, suggests that regions with stronger communication between certain molecules (e.g., COL1A1-CD36) are more vulnerable to developing the atrophy patterns seen in these conditions (FTLD subtypes). On the other hand, reduced interactions between genes with negative PLS weights such as LGI2-ADAM22/11 and FGF9-FGFR3/FLRT1 predispose to atrophy in the same conditions. Notably, cell-cell interactions from the same category as neuron-neuron can still be implicated in different functional processes, with some correlating positively or negatively with atrophy. Figure 3c displays surface plots (lateral views of the left hemisphere) illustrating the atrophy map for 4RTau alongside the interaction map for COL1A1-CD36. The analysis suggests that certain brain tissues with enhanced communication between COL1A1 and CD36 could be susceptible to develop atrophy specific to 4RTau.

Among the top 10% of pairs, 28 of 104 showed strong bi-directional interaction between astrocytes and neurons, along with prominent contributions from neuron-microglia (13 pairs) and neuron-neuron (24 pairs) (Fig. 3d). Overall, interactions involving both microglia (9 pairs) and astrocytes (9 pairs) were predominant in the first axis (see microglia-microglia, astrocyte-astrocyte, and microglia-endothelial), compared to endothelial-related interactions.

PANTHER database was used for gene enrichment since its collection of biological pathways is specific for cell signalling (Mi & Thomas, 2009). The top interacting molecules from the top 10% were significantly enriched for pathways including Slit/Robo-mediated axon guidance, Notch signalling, integrin signalling, and Alzheimer’s disease presenilin pathway (Fig.3e; p < 0.001, FDR-corrected).

### Second axis of cell-cell interactions explains tissue damage in PS1 and DLB

The second axis explained 7.61% of the covariance between interactions and atrophy patterns. Figure 4a shows the PLS weights for the top 5% of interacting pairs contributing to the second axis, with interactions such as FAM3C-GLRA2 (family with sequence similarity 3 and glycine receptor alpha 2) and LGI-related (leucine-rich glioma inactivated) interactions showing the largest contributions to explaining atrophy. The neurodegenerative conditions with positive weights (Fig. 4b) were PS1 mutation-related dementia (associated with EOAD), atrophy patterns in EOAD itself, and ALS. These results suggest that the atrophy patterns common to these three conditions were mostly explained by increased communication between LGI-related interactions, such as LGI3-ADAM22 and LGI2-ADAM23, and decreased interactions between pairs like FAM3C and GLRA2.

**Figure 4.**
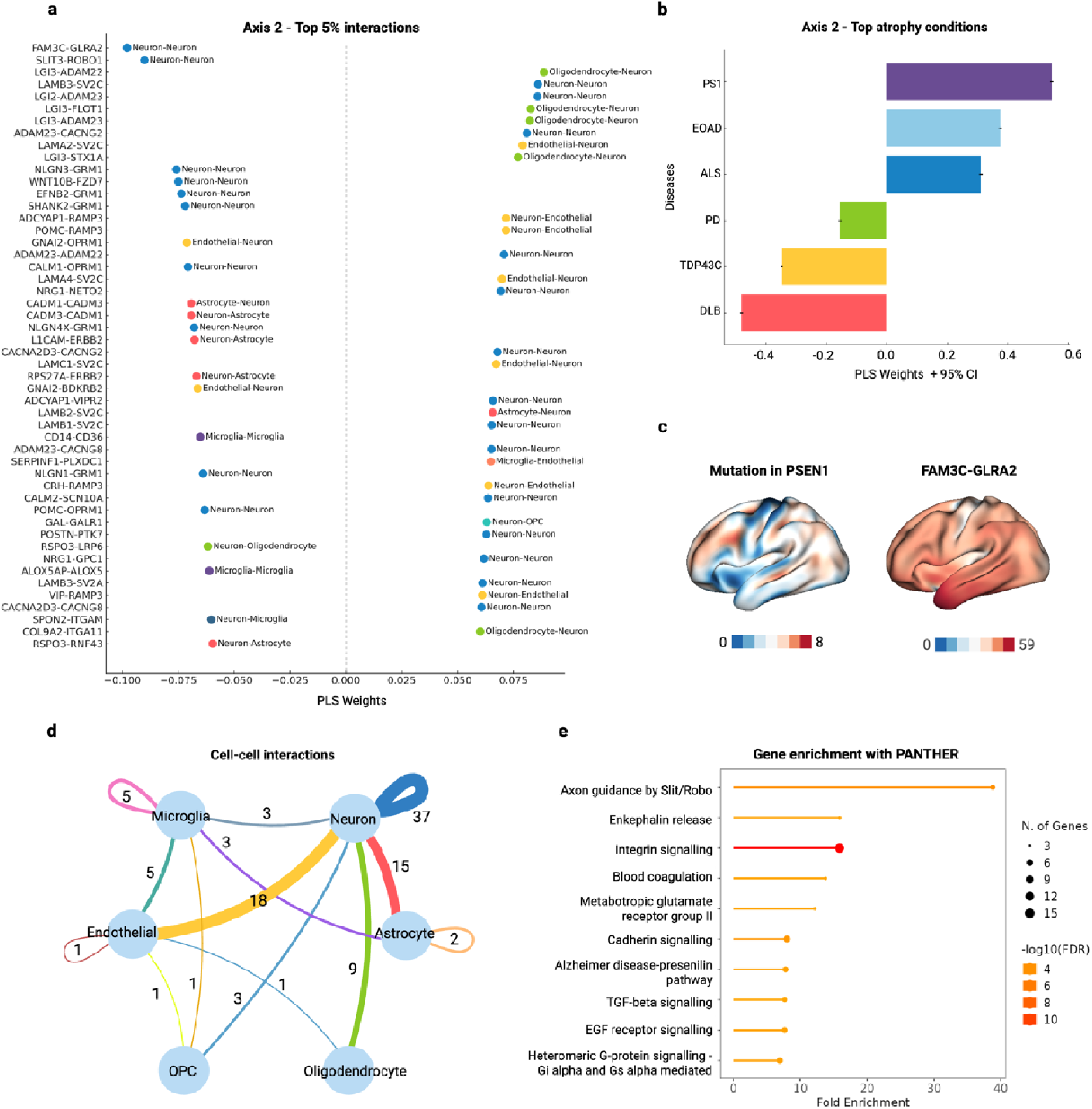
| Top interactions and neurodegenerative conditions contributing to the second axis. (**a**) PLS weights for the top 5% contributing interactions, colored by annotated cell-cell communication type. FAM3C-GLRA2 showed the highest contributions with negative weights. (**b**) Top neurodegenerative conditions with atrophy patterns contributing to the second axis. Mutation in PS1 and DLB showed the largest positive and negative contributions, respectively (error bars defined by 95% CI). (**c**) Surface plots (lateral views of the left hemisphere) displaying the atrophy map for PS1-related dementia and the LR interaction map for FAM3C-GLRA2. Color bars indicate the strength of cell-cell communication and tissue loss, with negative values indicating tissue enlargement relative to healthy controls. (**d**) Network visualization of cell-cell interactions counts from the top 10% annotated LR pairs, highlighting bi-directional signaling between neurons and endothelial cells, neurons and astrocytes, neurons and oligodendrocytes, and neurons with each other. Numbers on edges indicate counts of unique interactions. (**e**) Gene enrichment analysis using the PANTHER database, indicating significant enrichment of pathways related to Slit/Robo-mediated axon guidance, enkephalin release, integrin signalling, and Alzheimer’s disease presenilin (p < 0.001 FDR corrected).

Figure 4c displays the surface plots of the atrophy map for PS1 mutation carriers and the interaction map for FAM3C-GLRA2, highlighting their negative association in strength patterns. Genetic mutations in APP, such as PS1, modify Aβ biosynthesis and cause an autosomal dominant form of AD (George-Hyslop & Petit, 2005). Like PS1, FAM3C is a γ-secretase complex-binding protein. Previous studies have shown that FAM3C expression declines with aging and promotes Aβ accumulation in AD through its interaction with presenilin, with reduced FAM3C levels linked to the onset of sporadic AD, highlighting it as a potential therapeutic target (Hasegawa et al., 2014; Nishimura et al., 2020). Our results further support this association, as decreased FAM3C-GLRA2 communication indicates vulnerabilities to atrophy in PS1 mutation-related dementia and EOAD. Similarly, increased communication in FAM3C-GLRA2 and SLIT3-ROBO, along with decreased LGI-related interactions, were associated with the atrophy patterns common to DLB, TDP43 proteinopathy type C, and PD.

The interaction counts from the top 10% of pairs with the highest PLS weights captured a strong influence from neuron-neuron interactions (37 pairs), along with significant contributions from neuron-endothelial (18 pairs) and neuron-astrocyte (15 pairs) bidirectional interactions, as shown in Figure 4d. Unlike in the first axis, interactions involving neurons and oligodendrocytes, as well as neurons and endothelial cells, were more prominently represented in the second axis. For example, the LGI3 ligand, which showed multiple high contributions, is known to be uniquely secreted from oligodendrocytes in the brain and is involved in neuron-glial interaction processes, such as myelination (Miyazaki et al., 2024).

Gene enrichment analysis using the PANTHER database (Fig. 4e) indicated significant enrichment of pathways related to Slit/Robo-mediated axon guidance, enkephalin release, integrin signaling, and blood coagulation.

### Third axis of cell-cell interactions explains tissue atrophy patterns in PD and AD

The third axis explained 3.92% of the covariance (Fig. 5a), with prominent contributions from NPTX1-NPTXR and MET-related receptor interactions, also known as hepatocyte growth factor receptor, emphasizing their association with neurodegenerative atrophy patterns. Interactions related to brain-derived neurotrophic factor (BDNF) were also prominent, as BDNF regulates synaptic plasticity and cognitive functions and has been linked with AD (Gao et al., 2022-01-28; Ibrahim et al., 2022).

**Figure 5.**
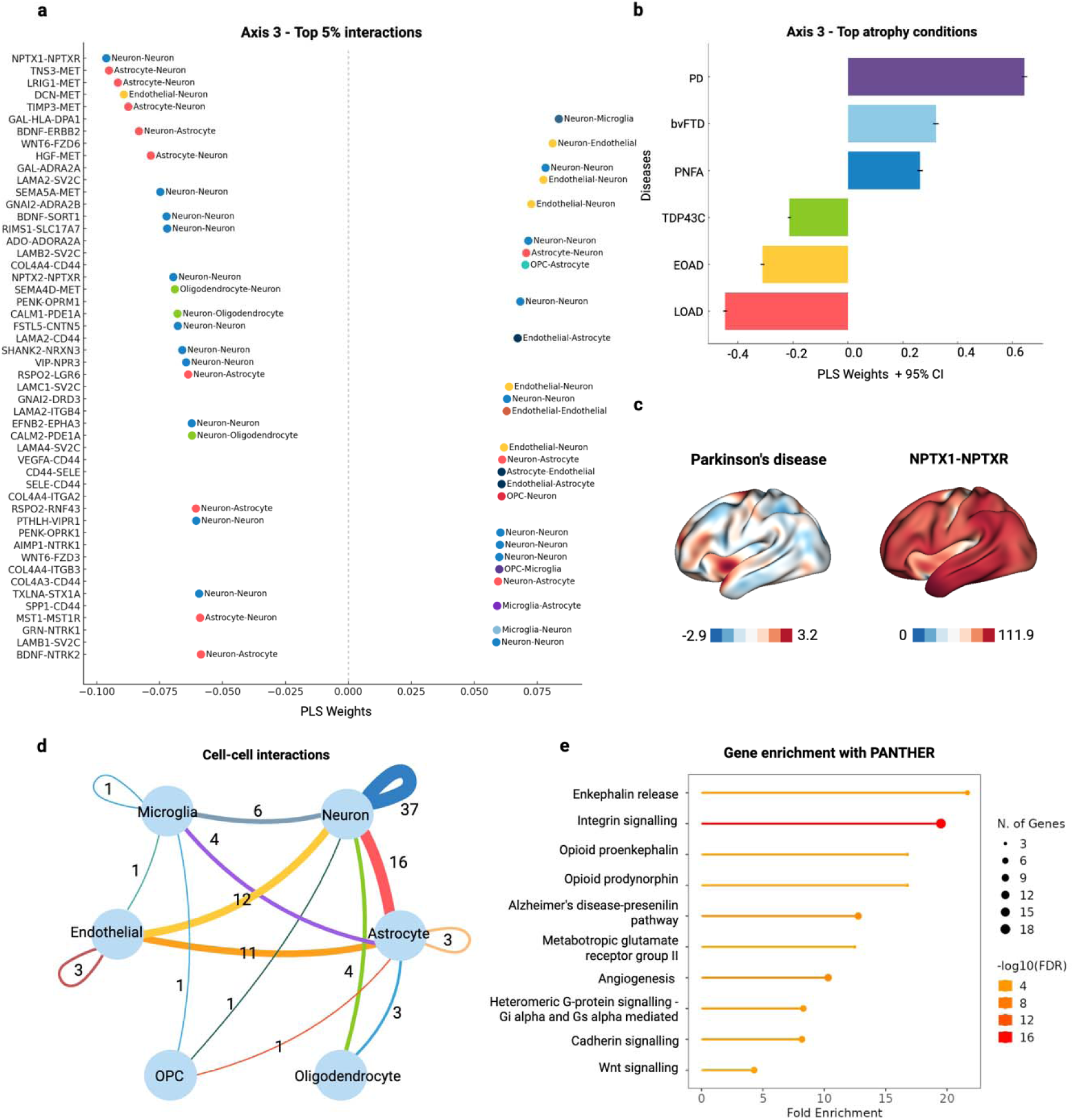
| Top interactions and neurodegenerative conditions contributing to the third axis. (**a**) PLS weights for the top 5% contributing interactions, colored by annotated cell-cell communication type, with NPTX1-NPTXR showing the highest contributions with negative weights. (**b**) Top six neurodegenerative conditions with atrophy patterns explaining the third axis the most. PD showed the largest contributions with positive weights, followed by LOAD and EOAD with negative weights (error bars defined by 95% CI). (**c**) Surface plots (lateral views of the left hemisphere) displaying the atrophy map for PD and the LR interaction NPTX1-NPTXR. Color bars indicate the strength of cell-cell communication and tissue loss, with negative values indicating tissue enlargement relative to healthy controls. (**d**) Network visualization of cell-cell interactions counts from the top 10% annotated LR pairs, pointing to strong bi-directional signaling between neuron-neurons and neuron-astrocytes, supporting by neuron-endothelial and endothelial-astrocyte. Numbers on edges indicate counts of unique interactions. (**e**) Gene enrichment with the PANTHER database, indicated significant enrichment of pathways related to enkephalin release, integrin signalling, opioid proenkephalin, and opioid prodynorphin (p < 0.001 FDR corrected).

Among the top six conditions contributing to the third axis (Fig. 5b), PD and bvFTD showed the largest positive weights, while LOAD and EOAD showed negative associations. These results suggest that decreased NPTX1-NPTXR and MET-related interactions may explain the atrophy patterns common to PD, bvFTD, and PNFA, while negatively associating with the atrophy observed in LOAD and EOAD.

Figure 5c illustrates brain surface maps for atrophy in PD and the spatial interaction map for NPTX1-NPTXR. Neuronal pentraxin 1 has previously been reported to be dysregulated in the brains of patients with PD (Warth Perez Arias et al., 2023). NPTX2 was also present among the top 20 contributing interactions to the third axis. Neuronal pentraxin II is known to be a cerebrospinal fluid biomarker of synaptic, motor, and cognitive dysfunctions in PD (Gómez de San José et al., 2021; Nilsson et al., 2023). Notably, GNAI2-DRD2 and GNAI2-DRD3 interactions, which showed strong communication across dopamine-enriched regions in Figure 2, including the substantia nigra involved in PD, were also present in the top 10% of the third axis.

Neuron-neuron interactions predominantly contributed to the third axis’ patterns (37 pairs), followed by 16 neuron-astrocyte bidirectional interactions (Fig. 5d). Neuron-endothelial and endothelial-astrocyte interactions also showed significant presence (12 and 11 pairs, respectively). The contribution from endothelial-astrocyte interactions distinguished the third axis from other axes. Gene enrichment analysis with the PANTHER database (Fig. 5e) highlighted pathways enriched in third axis. With an FDR-corrected p-value of 0.001, pathways such as enkephalin release, integrin signalling, opioid proenkephalin, and opioid prodynorphin were enriched.

### Identified cell-cell interactions in LOAD are reproduced in an independent population

Next, to validate our key findings beyond the used *in-silico* approach, we investigated whether the cell-cell interactions identified in neurotypical brains in the context of vulnerability remain relevant in the context of active neurodegeneration. Our initial analyses utilized AHBA data from non-diseased neurotypical brains in conjunction with atrophy measurements from individuals with neurodegenerative conditions. This approach raises an important question: to what extent are the identified neurotypical cell-cell interaction patterns, and their associations with regional vulnerability, preserved during actual disease progression? To address this question, we conducted complementary validation analyses using matched transcriptomic and atrophy data from the DLPFC of 375 individuals with LOAD from HBTRC (Zhang et al., 2013).

Atrophy measurements for the prefrontal cortex were available for all participants (Zhang et al., 2013), which enabled us to apply PLS correlation using the 1003 evaluated co-expression/interaction available pairs as observations, with variations in atrophy levels as the response variable. We then run PLS correlation using 1003 equivalent interactions derived from whole-brain transcriptomic AHBA data from healthy individuals and the whole-brain atrophy map from LOAD patients. This approach was an equivalent for the main results that we applied to get the three main axes, but for LOAD only.

We analyzed the top 10% (100 pairs) from each dataset and identified 14 overlapping molecular interactions (for all PLS results, see **Supplementary Table 5**). Figure 6a lists the top 10 interactions explaining atrophy in LOAD, where LGI2-ADAM11 can be seen prominent in both datasets. Figure 6b illustrates the distribution of weights for 14 overlapping interactions across both datasets. Variations in positive and negative values across axes result from certain gene expression data in the HBTRC dataset being negative after correction for covariates such as sex, age, and post-mortem interval, while in AHBA-LOAD atrophy values were indicated as either positive (greater atrophy) or negative (tissue enlargement compared to controls). Nonetheless, most interactions with the same interaction labels as Astrocyte-Astrocyte, Astrocyte-Neuron and Neuron-Neuron clustered together across both datasets.

**Figure 6.**
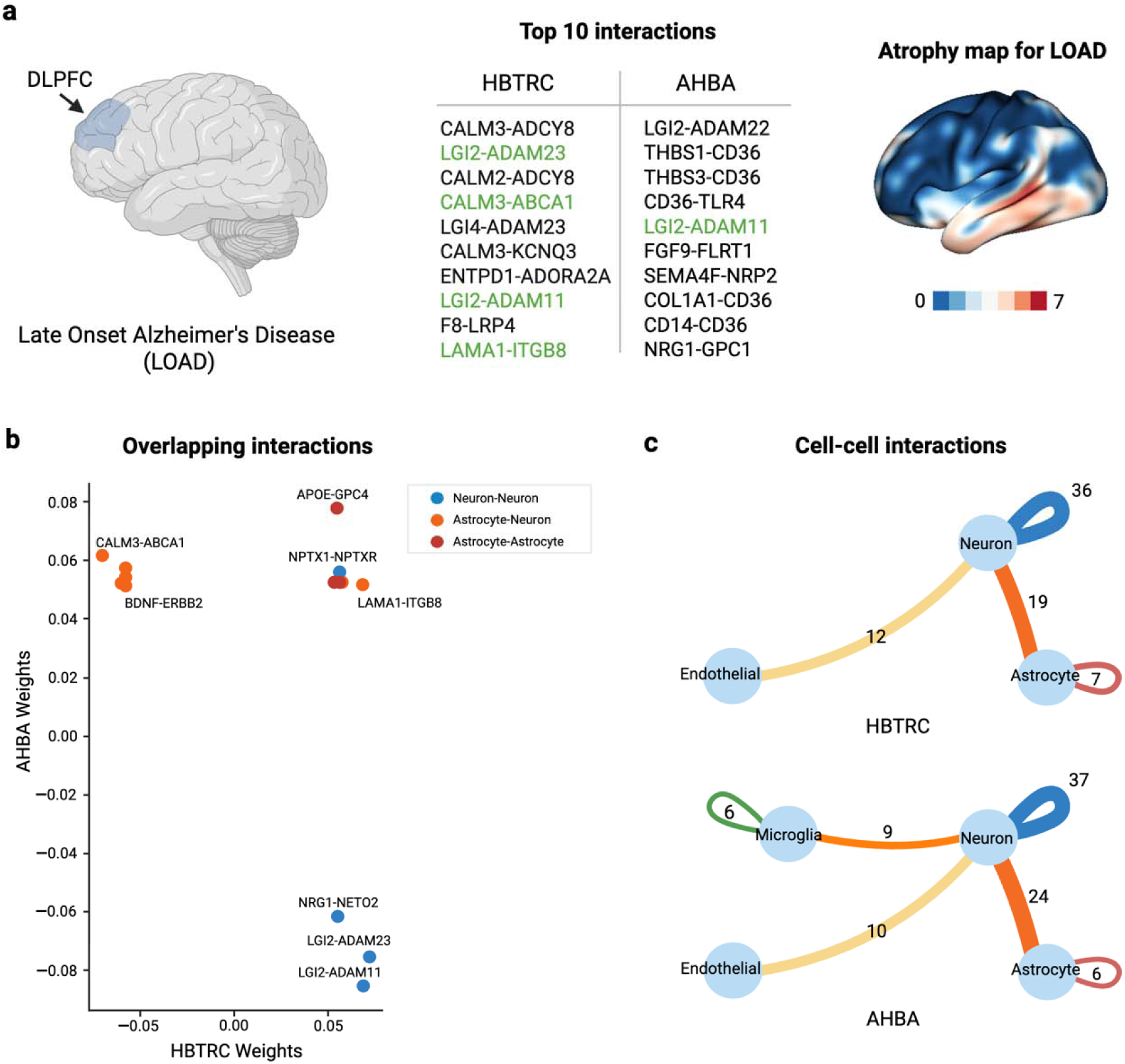
| Cross-validation of interactions linked to atrophy in LOAD. (**a**) Top 10 interactions contributing to atrophy in the DLPFC of patients with LOAD (left) from the HBTRC dataset and in whole healthy brains (right) from the AHBA, spatially aligned with atrophy maps of LOAD. Overlapping interactions between two datasets are highlighted in green. (**b**) Fourteen overlapping interactions found in the top 10% of interactions across both datasets, along with their corresponding PLS correlation weights. Pairs with the same interaction label grouped together. (**c**) Cell-cell interactions (with counts >5) present in the top 10% of pairs in both the HBTRC and AHBA datasets.

Figure 6c details the cell-cell interaction types most frequently occurring in the top 10% from both datasets. Both HBTRC and AHBA analyses demonstrated similar patterns, with neuron-neuron interactions being most frequent, followed by neuron-astrocyte and neuron-endothelial interactions. These results are different from the first axis of tissue vulnerability in several conditions (Fig. 3), where most of the interactions were captured between astrocytes and neurons, with only a few interactions between neurons and endothelial. Additionally, the AHBA-LOAD approach captured more neuron-microglia interactions (9 pairs) compared to the HBTRC (4 pairs).

Interactions between LGI2 (leucine-rich glioma inactivated 2) and the ADAM (a-disintegrin-and-metalloproteinase) family of neuronal proteins were the most prominent, explaining the association with atrophy in patients with LOAD and vulnerability to developing atrophy in LOAD. Interestingly, LGI2-ADAM22 and LGI2-ADAM11 were also in the top 10 contributing interactions to the first axis (Fig. 3). These interactions have been previously linked to epilepsy, synaptic transmission, and myelination (Kegel et al., 2013; Kozar-Gillan et al., 2023/04/03; Seppälä et al., 2011).

Other prominent overlapping interactions mentioned earlier were APOE-GPC4 from the first axis, which drives tau hypophosphorylation (Saroja et al., 2022), and NPTX1-NPTXR, which was the number one top interaction explaining atrophy in the third vulnerability axis. NPTX1 is known to play a critical role in synapse loss, neurite damage, and neuronal death induced by Aβ in LOAD (Abad et al., 2006). Additional pairs such as BDNF-ERB2 (also contributes to the third axis) and CALM2/3-ABCA1 have also been previously linked to neurodegeneration, with reduced BDNF levels contributing to the selectivity in neuronal degeneration in AD (Ibrahim et al., 2022) and ABCA1 regulating APOE lipidation, which affects Aβ aggregation and clearance in AD (Lewandowski et al., 2022). Overall, these results support that the cell-cell interactions associated with tissue vulnerability according to our *in-silico* approach, are also relevant in a large LOAD population.

### Comparison with interactions derived from snRNA-seq data

We assessed how spatial variation of interactions derived from bulk AHBA data across brain regions correlate with interactions derived from snRNA-seq data from the ABC atlas (Siletti et al., 2023). The ABC atlas consists of snRNA-seq measurements from 109 regions of neurotypical human donors (Siletti et al., 2023). Interaction strengths were measured across 76 regions defined by the Allen Human Reference Atlas (Ding et al., 2016), matching those available in the ABC dataset. For ABC, gene expression was calculated specifically from the cell types corresponding to the labelled ligands and receptors that we annotated for 1,037 pairs. For instance, if a gene was annotated as belonging to a specific cell type, only the expression of this gene from that particular cell type in the ABC atlas was used to calculate the cell-cell communication score. We conducted partial Spearman correlations with 500 permutations and FDR correction. Out of 1,027 LR pairs with available genes in ABC, 409 interactions (39.82%) showed significant correlation across both datasets.

Of the top 10% (104 interactions) contributing to the first axis, 43 interactions showed significant alignment (positive correlation) with their snRNA-seq counterpart across regions. Figure 7a illustrates regional differences in communication scores between bulk and snRNA-seq datasets. Difference values were normalized per region, with positive values indicating higher communication in AHBA interactions and negative values indicating higher expression in ABC interactions. Regions such as pons and septal nuclei (SEP) demonstrated larger differences between the two datasets compared to other regions. APOE-GPC5, SLIT-ROBO1/2, and NCAN-CDH2 demonstrated visibly higher communication in ABC interactions across many regions. Figure 7b depicts Spearman correlation coefficients (R) before permutation and FDR correction for these 43 interactions. Such pairs as SLIT-ROBO2 (R = 0.65) and PAPLN-SIRPA (R = 0.60) showed the strongest correspondence with the ABC validation set.

**Figure 7.**
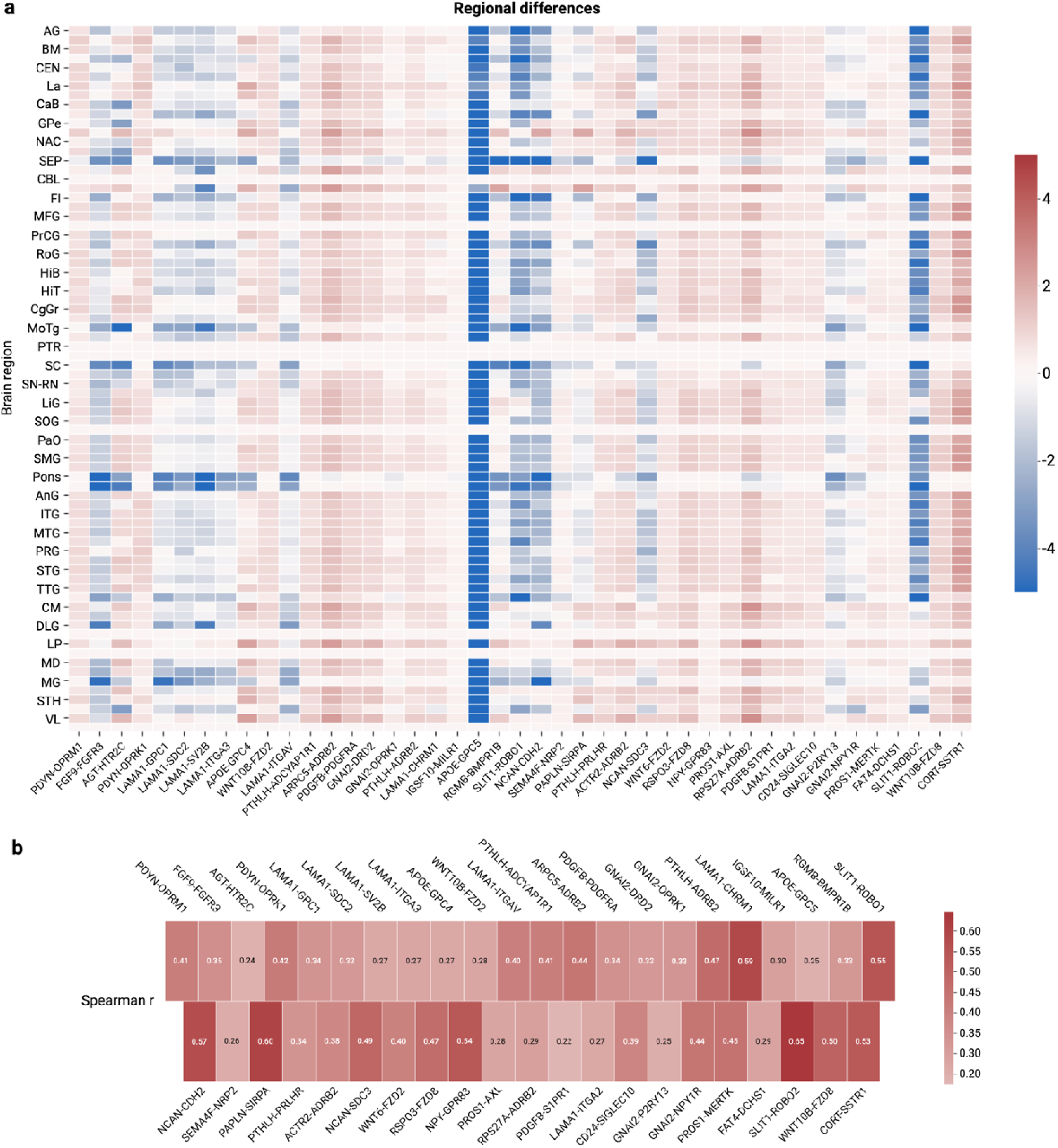
| Validation of bulk-derived cell-cell interactions with sn RNA-seq data from the ABC atlas. (**a**) Heatmap illustrating regional differences in communication between bulk-derived interactions (AHBA dataset) and interactions derived from snRNA-seq data (ABC atlas). Displayed interactions (n = 43) represent the subset among the top 10% of interactions contributing to the first axis that also showed significant positive partial correlation with the snRNA-seq counterpart. Values were normalized for each brain region, with positive values (red) indicating higher expression in bulk data and negative values (blue) indicating higher expression in ABC data. After normalization, zero values were replaced with epsilon (light pink colour) for better visualization. Regions (n = 76) were defined by the Allen Human Reference atlas. (**b**) Heatmap showing Spearman correlation coefficients (R) for these 43 significant interactions. Darker shades of red indicate stronger positive correlations, demonstrating consistent spatial variation between bulk and snRNA-seq data across brain regions.

### Disorder commonalities across the axes of cell-cell interactions

Lastly, we investigated commonalities across the 13 conditions in terms of their contributions to the major cell-cell interaction axes. Figure 8 presents a scatter plot of the normalized PLS loadings for the 13 disorders in the 3D space, using axes 1, 2, and 3 as coordinates. Based on the Silhouette criterion applied to the K-Means algorithm, four optimal clusters of disorders were identified.

**Figure 8.**
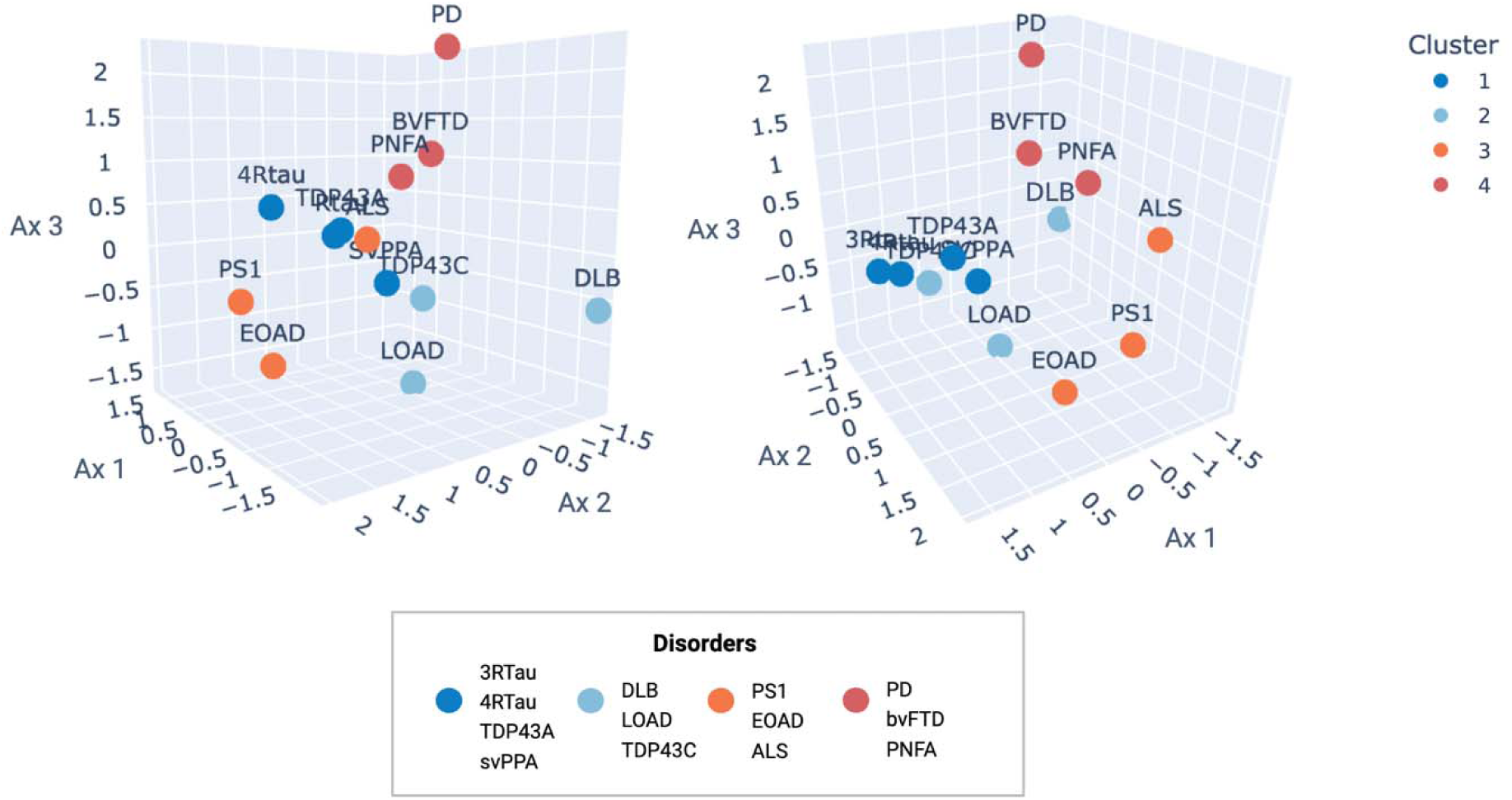
| Three-dimensional representation of PLS loadings across the neurodegeneration spectrum. Scatter plot illustrating the Z-score normalized PLS loadings for 13 neurodegenerative disorders in a 3D space, using axes 1, 2, and 3 as coordinates. The same plot is shown from two viewing angles to better visualize the spatial distribution of disorders. Based on Silhouette scores, K-Means clustering identified four optimal clusters: FTLD-related conditions (dark blue); TDP43C pathology, LOAD, and DLB (light blue); EOAD and ALS linked with PS1 mutations (orange); and bvFTD, PNFA, and PD (red).

In the first cluster (dark blue), FTLD-related conditions, including 3-repeat and 4-repeat tauopathies, TDP43A pathology, and svPPA, were grouped together. Interestingly, TDP43C pathology and LOAD clustered together with DLB in the second cluster (light blue). Atrophy patterns in TDP43C explained cell-cell interactions associated with all three axes. Unlike TDP43A that demonstrate similar patterns of frontotemporal atrophy to tauopathies, TDP43C involves greater left anterior-temporal tissue loss (Harper et al., 2017). Other FTLD conditions, such as the clinical subtypes bvFTD and PNFA, formed a separate cluster along with PD (red). Notably, both PD and DLB, which were positioned further away from the other conditions, are known synucleinopathies, unlike other conditions in this analysis that are often characterized by mixed pathologies (TDP-43, Aβ and tau).

Mutations in the PS1 clustered with EOAD and ALS, as these conditions exhibited the highest positive weights in the second axis. This finding is particularly interesting given the close relationship between EOAD and PS1 mutations, as PS1 mutations are the most common cause of autosomal dominant EOAD (Hutton & Hardy, 1997).

## Discussion

The systematic characterization of cell-cell communication across the brain and multiple neurological conditions remains technically challenging. To tackle this gap, here we reconstructed whole-brain maps of more than 1,000 molecular interactions specific to major brain cell types and conducted multimodal analysis to determine how cell-cell interactions account for spatial variation in tissue atrophy across neurodegenerative conditions. This *in-silico* multimodal approach uncovered axes of whole-brain cell-cell communication significantly explaining tissue vulnerability to neurodegeneration. Notably, our complementary validation analysis in an independent *post-mortem* LOAD cohort corroborated our initial findings. This convergence between our *in-silico* predictions and empirical observations in diseased tissues supports that the identified cellular communication networks represent crucial biological mechanisms underlying differential vulnerability across the neurodegenerative spectrum, rather than computational artifacts or coincidental associations. Together, these findings reveal the distinctive and shared cellular vulnerability across multiple neurodegenerative disorders, highlighting potential targets for therapeutic interventions.

Our results provide several key insights: (i) the first principal axis, explaining significant covariance with atrophy in subtypes of FTLD and AD, was dominated by astrocyte-neuron and astrocyte– and microglia-related interactions, such as FGF9-FGFR3 and CD36; ii) the second and third axes emphasized vascular endothelial-related interactions, effectively explaining atrophy patterns in mutations of PS1, DLB, AD, and PD, and featuring interactions like FAM3C-GLRA2, NPTX1-NPTXR, and MET; (iii) the enrichment analysis indicated such biological pathways as Slit/Robo-mediated axon guidance, enkephalin release, angiogenesis, integrin and cadherin signaling, and the Alzheimer’s disease presenilin pathway; and (iv) several critical interactions, such as LGI2-ADAM23/11, identified through our method to be relevant for LOAD tissue damage, were also found associating with atrophy in the DLPFC of patients with LOAD.

Despite the observed class imbalance, with neuron-neuron interactions being predominant in our analysis, bi-directional astrocyte-neuron crosstalk played a significant role in all of our findings, consistent with recent studies on cell-cell communication in AD (Bartas et al., 2024; Liu et al., 2024; Nam et al., 2023; Nanclares et al., 2021). For instance, a globally downregulated astrocyte-to-excitatory-neuron signaling axis was previously identified in the postmortem prefrontal cortex (PFC) of AD patients across two independent snRNA-seq datasets (Liu et al., 2024). Specifically, this study highlighted the downregulation of LR pairs involving AGT, APOE, PTN, and CALM ligands. Among the top 10% of pairs from the first axis, at least 14 interactions involved AGT, APOE, and several CALM-related interactions. Increased signalling from astrocytes to excitatory glutamatergic neurons was also observed in the entorhinal cortex of both patients with AD and 5xFAD mice (Bartas et al., 2024). Another snRNA study encompassing six brain regions in AD patients identified astrocytes, alongside both inhibitory and excitatory neurons, as exhibiting the most extensive gene expression changes across all regions (Mathys et al., 2024). These findings collectively align with our prior research showing that the cell abundance of astrocytes and microglia are strong predictors of regional atrophy across multiple neurodegenerative conditions (Pak et al., 2024).

The identified molecules, along with their associated cell types and signaling pathways, represent promising therapeutic targets for individual or multiple neurodegenerative conditions. The first axis included communication among several well-established AD risk genes, such as APOE, APP, APOC1, ABCA1, CD36, and TLR4. Importantly, previous studies investigating neuron-glial interactions in the PFC of AD patients across three independent snRNA-seq datasets have also identified APOE and APP among the most common interactions (Soelter et al., 2024). However, the role of several other identified interactions in our study has been limited or less established in the context of neurodegeneration and may need further investigation by the scientific community. For instance, molecules such as FGF9 and BDNF have previously been associated with atrophy and neuronal loss in neurodegeneration (Ibrahim et al., 2022; Thümmler et al., 2023). Molecules from the top interactions with the biggest contributions to the axes, such as COL1A1-CD36, FAM3C-GLRA2 and NPTX1-NPTXR, have been studied in neurodegeneration individually (Dobri et al., 2021; Gómez de San José et al., 2021; Nishimura et al., 2020), but their specific roles as interactive pairs influencing atrophy have not yet been explored. Finally, the interactions between LGI and ADAM families have been prominent in both axes of vulnerability and in independent dataset for LOAD from the HBTRC, yet their specific involvement in neurodegeneration has yet to be studied.

This study presents a number of limitations. We analyzed stereotypical atrophy patterns from diverse studies with varying protocols and methodologies. The interpolated gene expression values used for cell-cell interaction measures represent approximations that may smooth over important local cellular distinctions. The genes that we studied may be expressed across multiple cell types, suggesting cautious interpretation of cell-cell interaction annotations. To mitigate this concern, we included only genes consistently annotated across multiple gene marker databases. Our use of bulk RNA sequencing data limited resolution of cell-to-cell variability, and validation with snRNA-seq confirmed only 39.82% of interactions, suggesting that more comprehensive future validation with single-cell/single-nucleus sequencing may enhance cell-cell communication measure and cell-type annotation accuracy. The databases of LR pairs we employed showed significant class imbalance, with neuron-neuron interactions overrepresented compared to those involving non-neuronal cells. This potentially left important non-neuronal communication patterns, such as oligodendrocyte-related interactions, undetected. Moreover, the number of cell-cell interactions used in our analysis may not accurately reflect their true biological abundance; for instance, while we have more neuron-endothelial interactions, astrocyte-neuron interactions are likely more prevalent *in vivo*. Finally, most literature-based interpretations focused primarily on AD, underscoring the need for expanded investigation of identified interactions in other neurodegenerative conditions, particularly FTLD.

Our study also has several strengths. First, our method explores global spatial patterns of whole-brain cell-cell interactions linked to various conditions in a systematic way, providing a viable alternative until direct whole-brain sequencing of both healthy and diseased brains becomes feasible. Second, it correctly replicates known interactions in AD, FTD, and PD, as validated by existing literature, suggesting its capability in capturing biologically meaningful interactions. Third, our validation analysis demonstrated the biological accuracy of our reconstructed dopamine-related interaction maps, highlighting enhanced communication within established dopaminergic pathways and the correspondence with dopaminergic systems measured by PET/SPECT. Fourth, we validated specific LOAD-related interactions identified from healthy brains by demonstrating their alignment with atrophy patterns in the DLPFC from disease participants. Lastly, our study examined commonalities in molecular interaction patterns across distinct and related neurodegenerative conditions, where subtypes of the same condition (e.g., FTLD) or genetically related disorders (e.g., EOAD and PS1) showed similar patterns. Importantly, the generated maps of molecular interactions will be freely shared with the research community. Combined with the multimodal PLS tools (also available at our lab’s GitHub), these maps can serve as a scalable tool for investigating molecular basis of brain phenotypes and less-studied neurological conditions, such as psychiatric disorders, generating new hypotheses, and guiding experimental follow-up.

## Data availability

Gene expression data from the Allen Human Brain Atlas are publicly available at https://human.brain-map.org/static/download. Gene expression data from autopsied prefrontal cortex tissue of patients with late-onset Alzheimer’s disease, along with associated demographic and clinical information, are available from the Gene Expression Omnibus (GEO accession number: GSE44770, a subset of GSE44772). Atrophy maps for pathologically confirmed dementias can be accessed via NeuroVault (Collection ID: 1818) at https://identifiers.org/neurovault.collection:1818. Raw demographic and MRI data from Parkinson’s disease and amyotrophic lateral sclerosis (ALS) patients are available through the PPMI database (https://www.ppmi-info.org) and CALSNIC (http://calsnic.org; ClinicalTrials.gov Identifier: NCT02405182), respectively. Atrophy maps for clinical variants of frontotemporal dementia (FTD) are deposited on Zenodo at https://doi.org/10.5281/zenodo.10383492. Data from the FTLDNI initiative are available through the Laboratory of Neuroimaging (LONI) Image Data Archive (https://ida.loni.usc.edu). Curated databases of ligand–receptor interactions were retrieved from https://github.com/LewisLabUCSD/Ligand-Receptor-Pairs. MATLAB code for cross-correlational partial least squares (PLS) analysis is available at https://github.com/neuropm-lab/svd_pls. JuSpace toolbox was downloaded from https://github.com/juryxy/JuSpace. Over one thousand whole-brain maps of cell-cell interactions generated in this study will be made available on Zenodo upon publication. The braincellann R package for annotating brain-specific gene markers by cell type is available at https://github.com/nikaxpak/braincellann.

## Competing interests

The authors report no competing interests.

## Acknowledgments

This project was partly supported by the following awards to YIM: Canada Research Chair tier-2, NSERC Discovery grant, CIHR Project Grant 2020, Warren Y. Soper Charitable Trust, and Weston Family Foundation’s Transformational Research in AD 2020. VP was mainly supported by the Fonds de recherche du Québec (FRQ) doctoral scholarship and the Laszlo & Etelka Kollar Fellowship from the Faculty of Medicine and Health Sciences at McGill University and partly supported by the HBHL’s Theme 1 Discovery fund 2022–2025. We used the computational infrastructure of the McConnell Brain Imaging Center, supported in part by the *Brain Canada Foundation*.

## Notes

### Competing Interest Statement

The authors have declared no competing interest.

### Summary of Updates

Text throughout the manuscript has been improved for clarity and readability. We additionally expanded our analysis to include spatial correlations between dopamine-related cell-cell interaction maps and PET-derived measures of dopamine, providing validation of the transcriptomically-inferred dopamine signaling patterns.

